# Dihydropyrimidinase protects from DNA replication stress caused by cytotoxic metabolites

**DOI:** 10.1101/420893

**Authors:** Jihane Basbous, Antoine Aze, Laurent Chaloin, Rana Lebdy, Dana Hodroj, Cyril Ribeyre, Marion Larroque, Caitlin Shepard, Baek Kim, Alain Pruvost, Jérôme Moreaux, Domenico Maiorano, Marcel Mechali, Angelos Constantinou

**Affiliations:** Institute of Human Genetics (IGH), Univ Montpellier, CNRS, Montpellier, France.; Institut de Recherche en Infectiologie de Montpellier, CNRS, Université de Montpellier, Montpellier, France; Cancer Research Center of Toulouse (CRCT), Toulouse, France.; Institut du Cancer de Montpellier (ICM), Montpellier, France.; Center for Drug Discovery, Department of Pediatrics, Emory School of Medicine, Atlanta, GA 30322, USA.; Service de Pharmacologie et Immunoanalyse (SPI), Plateforme SMArt-MS, CEA, INRA, Université Paris-Saclay, Gif-sur-Yvette Cedex, France.

## Abstract

Imbalance in the level of the pyrimidine degradation products dihydrouracil and dihydrothymine is associated with cellular transformation and cancer progression. Dihydropyrimidines are degraded by dihydropyrimidinase (DHP), a zinc metalloenzyme that is upregulated in solid tumors but not in the corresponding normal tissues. How dihydropyrimidine metabolites affect cellular phenotypes remains elusive. Here we show that the suppression of DHP in cancer cell lines is cytotoxic. An increase in the level of dihydropyrimidines induced DNA replication and transcriptional stress. Cells lacking DHP accumulated DNA-protein crosslinks (DPCs), including covalently trapped DNA polymerase η. Furthermore, we show that the plant flavonoid dihydromyricetin inhibits human DHP activity. Cellular exposure to dihydromyricetin triggered DPCs-dependent DNA replication stress in cancer cells. This study defines dihydropyrimidines as potentially cytotoxic metabolites that may offer an opportunity for therapeutic-targeting of DHP activity in solid tumors.

## Introduction

The proliferation of cancer cells is associated with adjustments in metabolic activities required to satisfy high metabolic demands (Hanahan and Weinberg, 2011), and with frequent obstacles to the progression of replication forks that cause genomic instability (Hanahan and Weinberg, 2011; Macheret and Halazonetis, 2015). A variety of endogenous impediments result in the slowing or stalling of replication forks (Mirkin and Mirkin, 2007; Zeman and Cimprich, 2013). Activated oncogenes induce aberrant S phase entry without coordination with the production of limiting metabolites such as nucleotides (Bester et al., 2011). Furthermore, oncogenes allow the firing of replication origins within gene bodies, which induces replication/transcription conflicts (Macheret and Halazonetis, 2018). Specific DNA sequences can adopt non-canonical DNA structures that are difficult to replicate (Mirkin and Mirkin, 2007), such as poly (dA:dT) sequences that cause the breakage of replication intermediates at fragile sites under suboptimal DNA replication conditions (Tubbs et al., 2018). Misincorporated ribonucleotides (Huang et al., 2017; Lazzaro et al., 2012), DNA lesions caused by environmental agents and by reactive metabolic products such as reactive oxygen species and aldehydes (Langevin et al., 2011; Lindahl, 1993; Pontel et al., 2015) interfere with the progression of replication forks. Furthermore, cellular metabolites can yield structurally diverse DNA-protein crosslinks (DPCs) that precipitate the loss of cellular functions, including transcription and DNA replication (Tretyakova et al., 2015). Interference with the progression of replication forks is exploited therapeutically through stress overload by chemotherapeutic agents that induce DNA adducts to block DNA replication (Luo et al., 2009).

DNA replication stress induces the accumulation of 70 to 500 long stretches of single-stranded DNA (Hashimoto et al., 2010; Sogo et al., 2002; Zellweger et al., 2015), which trigger a protein kinase cascade orchestrated by the checkpoint kinase ATR and its effector kinase Chk1 (Guo et al., 2000; Hekmat-Nejad et al., 2000; Liu et al., 2000; Zhao and Piwnica-Worms, 2001). ATR signalling promotes cell and organismal survival through coordination of DNA repair and DNA replication with cell physiological processes including cell cycle progression and transcription (Ciccia and Elledge, 2010). ATR accumulates at DNA replication sites through recognition of RPA by its partner protein ATRIP (Zou and Elledge, 2003). ATR signalling limits the accumulation of single-stranded DNA (Toledo et al., 2013). Above a critical threshold of single-stranded DNA, exhaustion of nuclear RPA provokes catastrophic DNA breaks throughout the nucleus (Toledo et al., 2017; Toledo et al., 2013).

Accumulating evidence indicates that carcinogenesis is associated with alterations in the level of enzymes that degrade the pyrimidines uracil and thymine (Edwards et al., 2016; Naguib et al., 1985; Shaul et al., 2014; Wikoff et al., 2015). Whether and how rewiring of this pyrimidine catabolic pathway support tumors progression remains poorly defined. Uracil and thymine are degraded in three enzymatic steps (Figure 1A). First, dihydropyrimidine dehydrogenase (DPD) reduces the pyrimidine ring of uracil and thymine with hydrogen and yields 5, 6-dihydrouracil and 5, 6-dihydrothymine (dihydropyrimidines). Second, the saturated rings between position 3 and 4 are opened by dihydropyrimidinase (DHP). Third, β-ureidopropionase (BUP-1) degrades the β-ureidopropionic acid and β-ureidoisobutyric acid products formed by DHP into β-alanine and β-aminoisobutyric acid. DPD activity has been detected in all tissues examined, but the activity of DHP and BUP-1 is essentially restricted to the liver and the kidney (van Kuilenburg et al., 2006). In cancer cells, however, the level of pyrimidine degradation activities is considerably altered. Increased expression of DPD has been observed in human hepatocellular carcinoma (Yoo et al., 2009). Human skin cutaneous melanomas progressing towards metastatic tumors accumulate mutations in DPD and up-regulate the expression of the genes encoding DPD and DHP (Edwards et al., 2016). The accumulation of 5, 6-dihydrouracil is a distinct metabolic feature of early lung adenocarcinoma (Wikoff et al., 2015). Intriguingly, an increase in the concentration of dihydropyrimidines in epithelial breast cancer cells supports the acquisition of aggressive mesenchymal characteristics (Shaul et al., 2014). How dihydropyrimidines affect cellular phenotypes, however, remains elusive. Pioneering studies have identified dihydropyrimidinase (DHP) activity as a good marker of tumorigenicity and a target for cancer therapy (Naguib et al., 1985). Whereas hardly detectable in normal extrahepatic and kidney tissues (van Kuilenburg et al., 2006), the activity of DHP is strikingly high in human carcinomas of the lung, colon, pancreas, salivary gland and stomach (Naguib et al., 1985).

**Figure 1.**
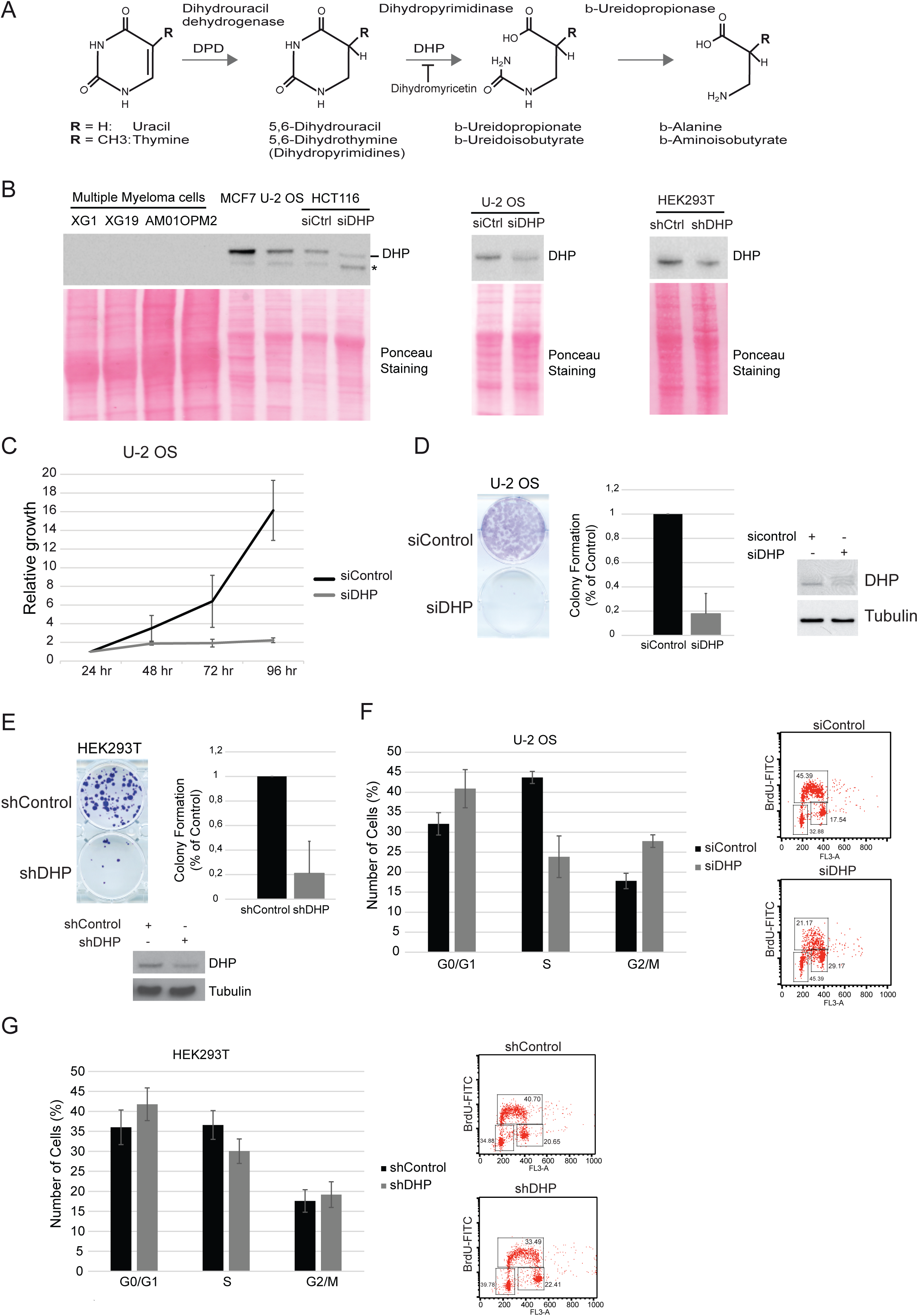
Dihydropyrimidinase depletion affects epithelial cancer cells proliferation. (A) Schematic representation of the pyrimidine degradation pathway. (B) DHP was probed by Western blotting in the indicated transformed cells. When indicated, DHP was knocked down using anti-DHP siRNA or shRNA molecules with distinct target sequences. Ponceau staining was used as loading control. * non specific signal. One representative experiment is shown from 3-6 biological replicates. (C) U-2 OS cells were transfected with control or anti-DHP siRNA (siDHP) and their viability was assessed during four days using the MTT cell growth assay. Mean viability is representative of three independent biological replicates. Error bars represent +/-S.D. (D) Colony-forming assay of U-2 OS cells after transfection with control or anti-DHP siRNA (siDHP). A representative image is shown. An histogram represents the quantification of colony formation. Data shown are averages over three independent biological replicates with two technical replicates for each. Error bars represent +/-S.D. (E) HEK293T cells transfected with control or anti-DHP shRNA were analyzed as described in (E). (F) Histogram representing the percentage of U-2 OS cells, 72 hours after transfection with control or anti-DHP siRNA (siDHP) in G0/G1, S and G2/M phases. Data shown are averages over three independent biological replicates. Error bars represent +/-S.D. (G) Histogram representing the cell cycle distribution of HEK293T cells 72 hours after transfection with control or anti-DHP shRNA Data shown are averages over three independent biological replicates. Error bars represent +/-S.D.

Here we show that suppression of DHP in cancer cell lines induces DNA replication stress, as revealed by the accumulation of single-stranded DNA, by the induction of ATR/Chk1 signaling and by the slowing of replication fork progression. Depletion of DHP also attenuates transcription activity, stabilizes p53 and eventually blocks cell proliferation. The addition of dihydropyrimidines to Xenopus egg-extracts induces the formation of abnormal DNA replication products. IN DHP depleted cells, DNA replication and transcriptional stress correlates with the accumulation of DNA-protein crosslinks (DPCs). Thus, we suggest that dihydropyrimidines yield DPCs that directly interferes with DNA-templated processes. We found that the flavonoid dihydromyricetin inhibits the activity of purified human dihydropyrimidinase. Addition of dihydromyricetin in the cell culture medium induces the accumulation of DNA-protein crosslinks and interferes with the progression of replication forks. These findings indicate that unless degraded by dihydropyimidinase, the amount of dihydropyrimidines produced in cancer cell cultures is sufficient to block DNA templated processes.

## Results

### Suppression of DHP induces DNA replication stress and inhibits cell proliferation

Dihydropyrimidinase (DHP) is mainly expressed in the liver and the kidney (van Kuilenburg et al., 2006). Yet, we detected DHP in the epithelial cancer cells MCF7 (breast adenocarcinoma), U-2 OS (osteosarcoma), HCT116 (colorectal carcinoma), and HEK293T (embryonic kidney) (Figure 1B). By contrast, DHP was not detectable in multiple myeloma cells (Figure 1B), consistent with the absence of DHP activity in hematopoietic malignant cells (Naguib et al., 1985). The knockdown of DHP by means of siRNA in HCT116, U-2 OS and HEK293T cells confirmed the specificity of the anti-DHP signal (Figure 1B).

As DHP is produced in different epithelial cancer cells, we set out to explore the phenotypic consequences of DHP depletion. Depletion of DHP in U-2 OS cells blocked cell proliferation measured by colorimetric and by colony forming assays (Figure 1C, D). The RAD51 mRNA transcript is sensitive to siRNA-mediated depletion (Adamson et al., 2012). To exclude potential off-target effects of siRNAs, we used a shRNA with a different target sequence to suppress DHP in HEK293T cells. The growth of HEK293T cells was severely compromised upon depletion of DHP with a shRNA (Figure 1E). These observations indicate that DHP is essential for the proliferation of these cancer cell lines. Thus, we could not generate *DPYS* (encodes DHP) knockouts cell lines by CRISPR/CAS9 genome editing to explore the function of DHP.

Flow cytometry cell cycle analyses (FACS) 72-hours post-transfection revealed alterations in the cell cycle distribution of osteosarcoma U-2 OS cells treated with an anti-DHP siRNA (Figure 1F). In comparison with control cells, suppression of DHP reduced the fraction of U-2 OS cells in S phase. The impact of DHP depletion was less pronounced in the transformed HEK293T cell line (Figure 1G), perhaps reflecting differences in the dependency on DHP activity or in the efficiency of DHP depletion. To evaluate further the capacity of DHP-depleted HEK293T cells to proceed through the cell cycle, we performed pulse-chase experiments with the nucleotide analogue BrdU. Cells were labeled with BrdU for 30 minutes and sorted by two-dimensional flow cytometry at the indicated time of incubation in BrdU free medium. After six-hours incubation in BrdU-free medium, the majority of control cells had completed S phase and a significant proportion proceeded to the G1 phase (9.27 %), whereas the proportion of DHP-depleted HEK293T cells to reach G1 was reduced by half (Figure S1A).

To explore the cause of the cell cycle delays, we analyzed DNA replication tracks at the single molecule level by DNA fiber labeling. Replication tracks were dually labeled with two consecutive pulses of fluorescent nucleotide analogues iodo-deoxyuridine (IdU) and chloro-deoxyuridine (CldU) for 30 minutes each. We assessed the progression of isolated replication forks by measuring the length of CldU tracks adjacent to IdU tracks. The knockdown of DHP by siRNA or shRNA molecules with different target sequences reduced the length of replication tracts in U-2 OS or HEK293T cells (Figure 2A and S1B). Re-expression of a siRNA-resistant myc-tagged DHP protein in DHP knockdown U-2 OS cells partially rescued replication progression (Figure 2A). This confirms that the DNA replication defect is a direct consequence of DHP depletion.

**Figure 2.**
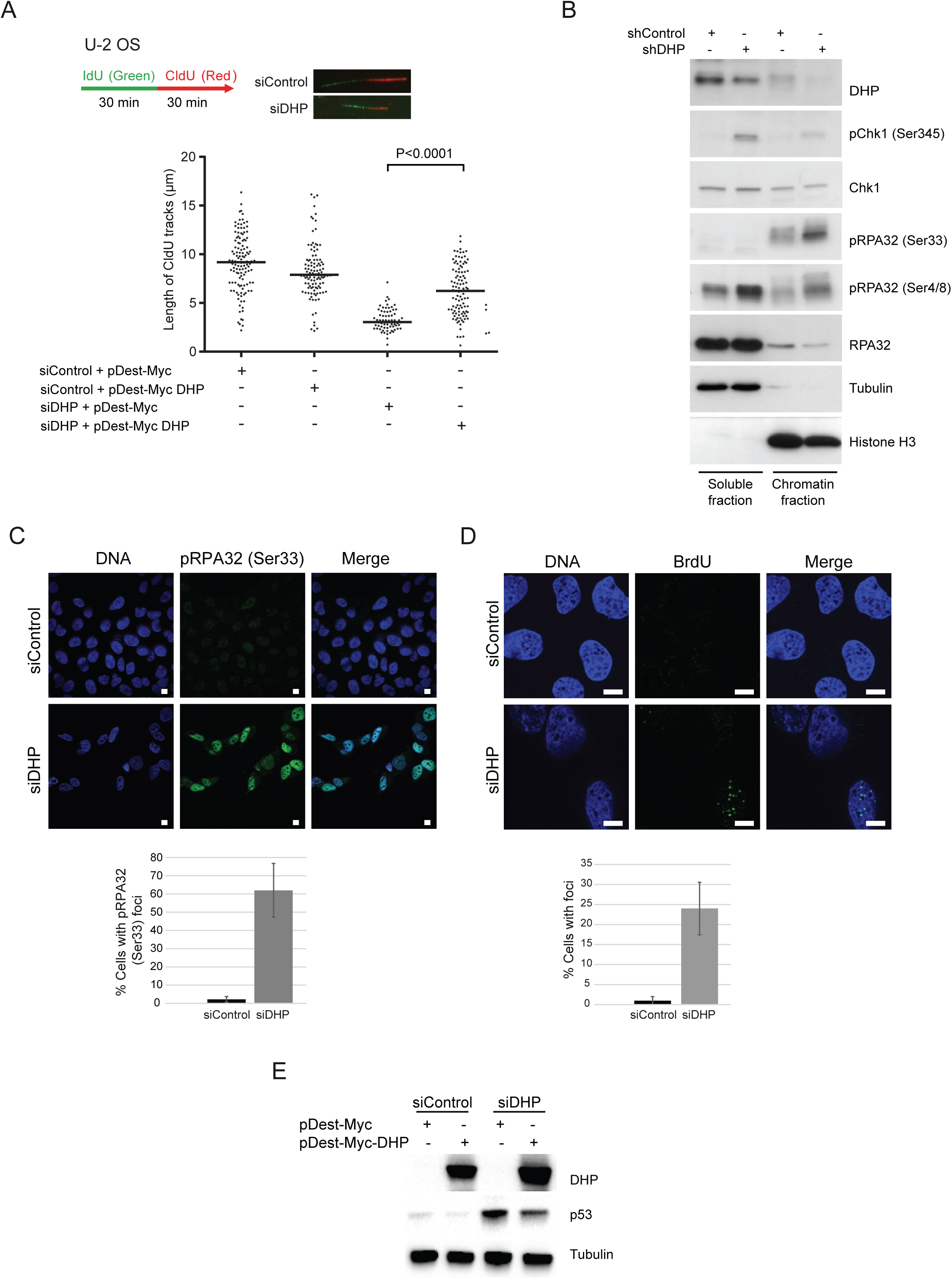
Suppression of DHP interferes with replication fork progression and induces activation of DNA damage responses. (A) Experimental scheme: 72 hours after transfection with control or anti-DHP siRNA (siDHP), cells were labeled with two consecutive pulses of 30 minutes with CldU and IdU, as indicated. DNA was streched out on glass slides and newly synthesized DNA was revealed by immunofluorescence. Graphic representation of replication track lengths in U-2 OS cells co-transfected with control or anti-DHP siRNA (siDHP) along with control plasmid or a plasmid encoding siRNA-resistant DHP. The bar dissecting the data points represents the median of 100 tracts length from one biological replicate. Differences between distributions were assessed with the Mann-Whitney rank sum test. (B) DHP knockdown HEK293T cells were subjected to subcellular fractionation and probed by western blotting with the indicated antibodies. *Tubulin and histone H3* were used as *loading controls* for the cytoplasmic and chromatin *fractions*, respectively. One representative experiment is shown from two biological replicates. (C) Immunofluorescence staining of Ser33 phospho RPA32 in control and DHP knockdown U-2 OS cells (siDHP). DNA was stained by Hoechst. Bars indicate 10 µm. Bottom panel: Histogram representing the percentage of Ser33 pRPA32 foci positive cells in a population of 100 cells. Data from three independent biological replicates are represented as mean +/-S.D. (100 cells were counted per experiment). (D) Control and DHP knockdown U-2 OS cells (siDHP) were uniformly labeled with BrdU before immunofluorescence staining in native conditions with an anti-BrdU antibody. DNA was stained by Hoechst. Bars indicate 10 µm. Right panel: Histogram representation of the percentage of ssDNA positive cells. Values are the mean ± SD of three independent biological replicates (100 cells were counted per experiment). (E) Western blotting analysis with the indicated antibodies of whole-cell extracts from control and DHP knockdown U-2 OS cells (siDHP) complemented or not with a siRNA-resistant DHP cDNA, as indicated. One representative experiment is shown from three biological replicates.

Next, we probed cells for indicators of DNA replication stress by monitoring ATR/Chk1 signaling. We observed spontaneous accumulation of Chk1 phosphorylated on Ser345 in the soluble fraction and of RPA32 phosphorylated on Ser 4/8 and Ser33 in the chromatin fraction of DHP-depleted HEK293T cells (Figure 2B). Phospho RPA32 (Ser33) also accumulated in HEK293T transfected cells with a different anti-DHP shRNA (Figure S1C). To confirm this observation, we visualized RPA foci and phospho RPA signals by means of immunofluorescence microscopy. RPA32 and phospho RPA32 (ser33) signals accumulated in U-2 OS cells transfected with anti-DHP siRNA (Figure S1D and 2C), and in HEK293T cells transfected with a distinct anti-DHP shRNA (Figure S1E). To verify that the formation of RPA32 foci correlates with the accumulation of single-stranded DNA (ssDNA), we probed ssDNA by immunofluorescence microscopy using uniform BrdU labeling and BrdU detection in native conditions (Raderschall et al., 1999). Nearly 30 % of DHP-depleted cells exhibited multiple and distinct BrdU signals indicative of severe replication-associated defects (Figure 2D). Last, we monitored the level of p53 that is stabilized in response to genotoxic stress. Depletion of DHP markedly increased the level of p53 in U-2 OS cells (Figure S1F). Expression of a siRNA-resistant cDNA encoding DHP reduced the level of p53, confirming that the stabilization of p53 results, at least in part, from the depletion of DHP (Figure 2E). Collectively, these data indicate that suppression of DHP induces DNA replication stress, at least in a subset of cancer cell lines.

### Accumulation of dihydropyrimidines induces DNA replication stress

Next, we sought to investigate how suppression of DHP inhibits fork progression. The rate of DNA chain elongation is dependent on the pool of available deoxyribonucleotides. Thus, we measured the impact of DHP depletion on dNTPs levels using a single nucleotide incorporation assay (Diamond et al., 2004). The level of dNTPs increases in proliferating cells and fluctuates during the cell cycle (Lane and Fan, 2015). Since the suppression of DHP has consequences on cell growth and cycle progression, cells lacking DHP may exhibit altered dNTPs levels. Consistent with this, the suppression of DHP in HEK293T and U-2 OS cells led to a reduction in the global level of dNTPs (Figure S2A and S2B). To test if alterations of dNTPs levels were responsible for the defect in fork progression observed in DHP-depleted cells, we complemented the cell culture medium with saturating concentrations of nucleosides and measured the length of CldU-labeled replication tracks from isolated replication forks. Addition of nucleosides in the cell culture medium markedly increased the length of replication tracks in shControl HEK293T cells (Figure S2C), with a median fold stimulation of 1.7x. This data indicates that nucleosides are limiting in these cells. By contrast, saturation of DHP-depleted cells with a cocktail of nucleosides did not markedly increase the length of replication tracks (Figure S2C). Therefore, changes in dNTPs levels are not the primary cause of replication stress in these cells. Consistent with this interpretation, addition of an excess of nucleosides in the cell culture medium of DHP-depleted cells did not attenuate the accumulation of p53 and the phosphorylation of RPA32 on Ser33 (Figure S2D). Measurements of the global pool of dNTPs does not give insights into local levels of dNTPs available to the DNA replication machinery (Techer et al., 2016).

The rate of replication fork progression is primarily determined by the amount of DNA damage and the level of activated p53, not by the global concentrations of dNTPs (Techer et al., 2016). Thus, we considered the possibility that DNA replication stress in DHP-depleted cells was caused by the accumulation of dihydropyrimidines. To test this, we measured the cellular concentration of uracil and its breakdown product dihydrouracil by liquid chromatography and mass spectrometry (LC-MS). DHP-depletion in U-2 OS cells yielded a four-fold increase in the molar ratio of intracellular dihydrouracil/uracil (Figure 3A). This indicate that the transient knockdown of DHP is sufficient to raise the intracellular concentration of dihydropyrimidines. To counteract the accumulation of dihydropyrimidines in DHP-knockdown cells, we co-depleted dihydropyrimidine dehydrogenase (DPD), the first enzyme in the pyrimidine catabolic pathway that produces dihydropyrimidines (Figure 1A). Suppression of DPD in DHP-depleted cells rescued the rate of fork progression to the level of control cells (Figure 3B). Consistent with this result, the levels of p53, of Ser33 phosphorylated RPA32 and the intensity of RPA32 foci were close to normal when both DPD and DHP enzymes were depleted (Figure 3C and 3D). Furthermore, exposure of U-2 OS cells for five minutes to 10 or 40 mM 5, 6-dihydrouracil in a hypotonic solution induced, 48 hours later, the accumulation of p53 and the phosphorylation of RPA32 on Ser33 (Figure 3E). Altogether, these observations suggest that the DNA replication stress phenotype of DHP-depleted cells is the consequence of the accumulation of dihydropyrimidines.

**Figure 3.**
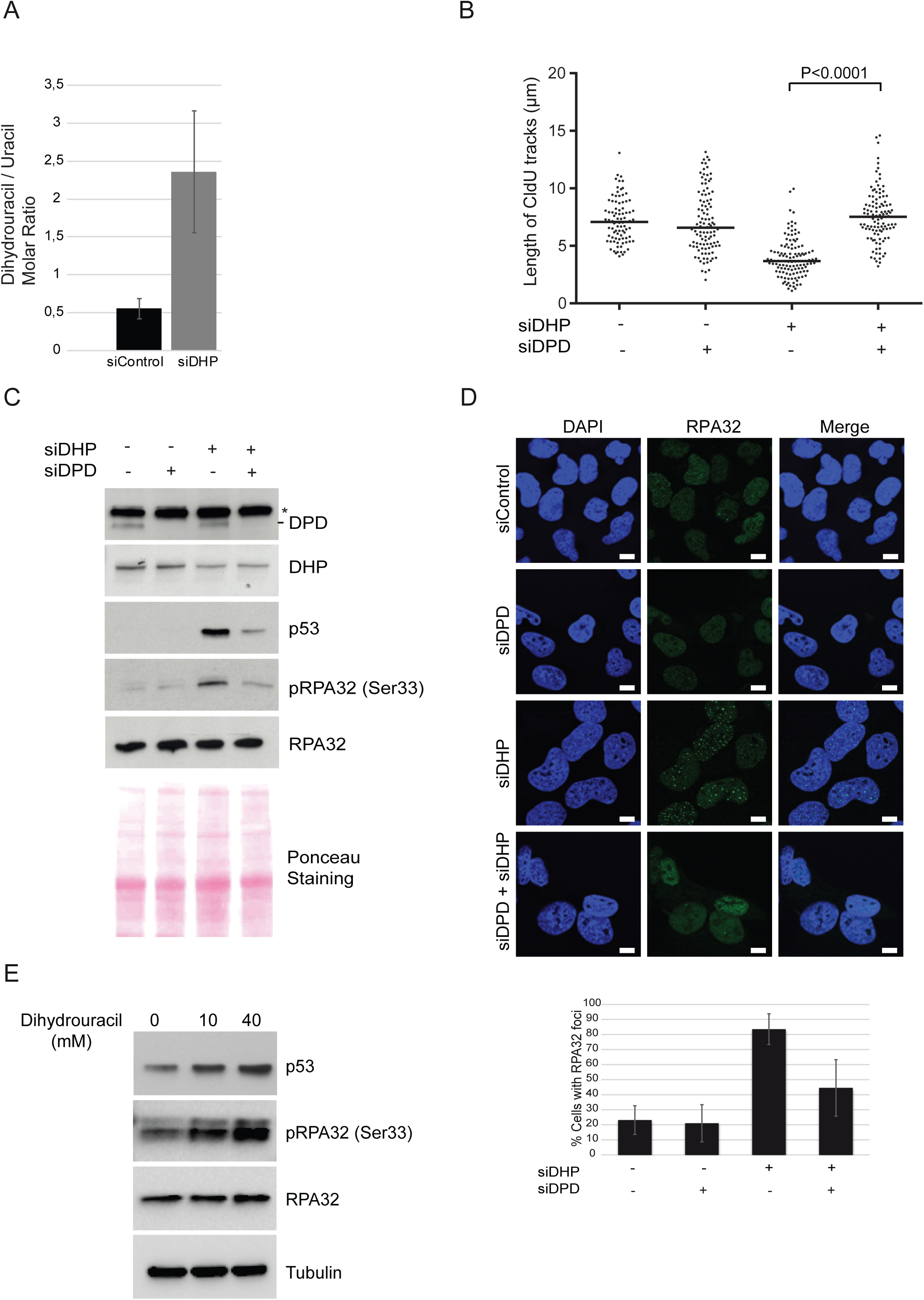
Dihydropyrimidines accumulation in transformed cells induce DNA replication stress. (A) The concentrations of dihydrouracil and uracil were measured in U-2 OS cells transfected with control or anti-DHP siRNA (siDHP). The ratio of molar concentrations between the two metabolites in each sample is presented. Data from three independent biological replicates, with three technical replicates for each, are represented as mean +/-S.E.M. (B) Replication tracks were labelled with two consecutive pulses of 30 minutes with CldU and IdU in U-2 OS cells transfected with the indicated siRNAs. Graphic representations of replication track lengths measured in μm (y axis). The bar dissecting the data points represents the median of 100 tracts length from one biological replicate. Differences between distributions were assessed with the Mann-Whitney rank sum test. (C) Western blot analysis with the indicated antibodies of whole cell extracts from U-2 OS transfected with anti-DHP and anti-DPD siRNAs, as indicated, Ponceau staining was used as control of protein loading and transfer. * non-specific band. One representative experiment is shown from two biological replicates. (D) RPA32 immunofluorescence staining of U-2 OS cells transfected with the indicated siRNAs. Bars indicate 10 µm. DNA was stained by Hoechst. Bottom panel: Histogram representation of the percentage of RPA32 foci-positive cells in a population of 100 cells. Data from three independent biological replicates are represented as mean +/-S.D. (E) U-2 OS cells were incubated for 5 min in a hypotonic buffer (50 mM KCl, 10 mM Hepes) containing 10 or 40 mM Dihydrouracil and resuspended into fresh cell culture medium for 48 hr prior to lysis and analysis by western blot with the indicated antibodies. One representative experiment is shown from two biological replicates.

### Dihydropyrimidine accumulation induces transcriptional stress

The data thus far indicate that dihydropyrimidine metabolites induce DNA replication stress. We reasoned that dihydropyrimidines could induce directly or indirectly the formation of DNA adducts and interfere with DNA-templated processes including transcription. Thus, we explored the effect of DHP loss on global transcription activity. Nascent RNA were pulse labelled for 20 minutes with the modified RNA precursor 5-ethynyluridine (EU) and overall transcription activity was evaluated via fluorescent-based quantification. In comparison with control cells, we observed a drop of EU incorporation in DHP-depleted U-2 OS cells (Figure 4A). Next, we used the anti-RNA/DNA hybrids S9.6 antibody to visualize R-loops by immunofluorescence staining. R-loops are induced by defects in the processing of nascent pre-mRNAs or by the accumulation of negative supercoiling behind RNA polymerases (Li and Manley, 2005; Tuduri et al., 2009). Immunofluorescence staining experiments revealed a significant increase in the level of nuclear RNA/DNA hybrids in DHP-depleted cells compared to control cells after excluding nucleolar signals from the analysis (Figure 4B). R-loops and nucleolin immunofluorescence staining revealed alterations in the morphology of nucleoli, which appeared more condensed and rounded (Figure 4B). These observations indicate that the accumulation of dihydropyrimidine metabolites in DHP-depleted cells induces transcriptional stress.

**Figure 4.**
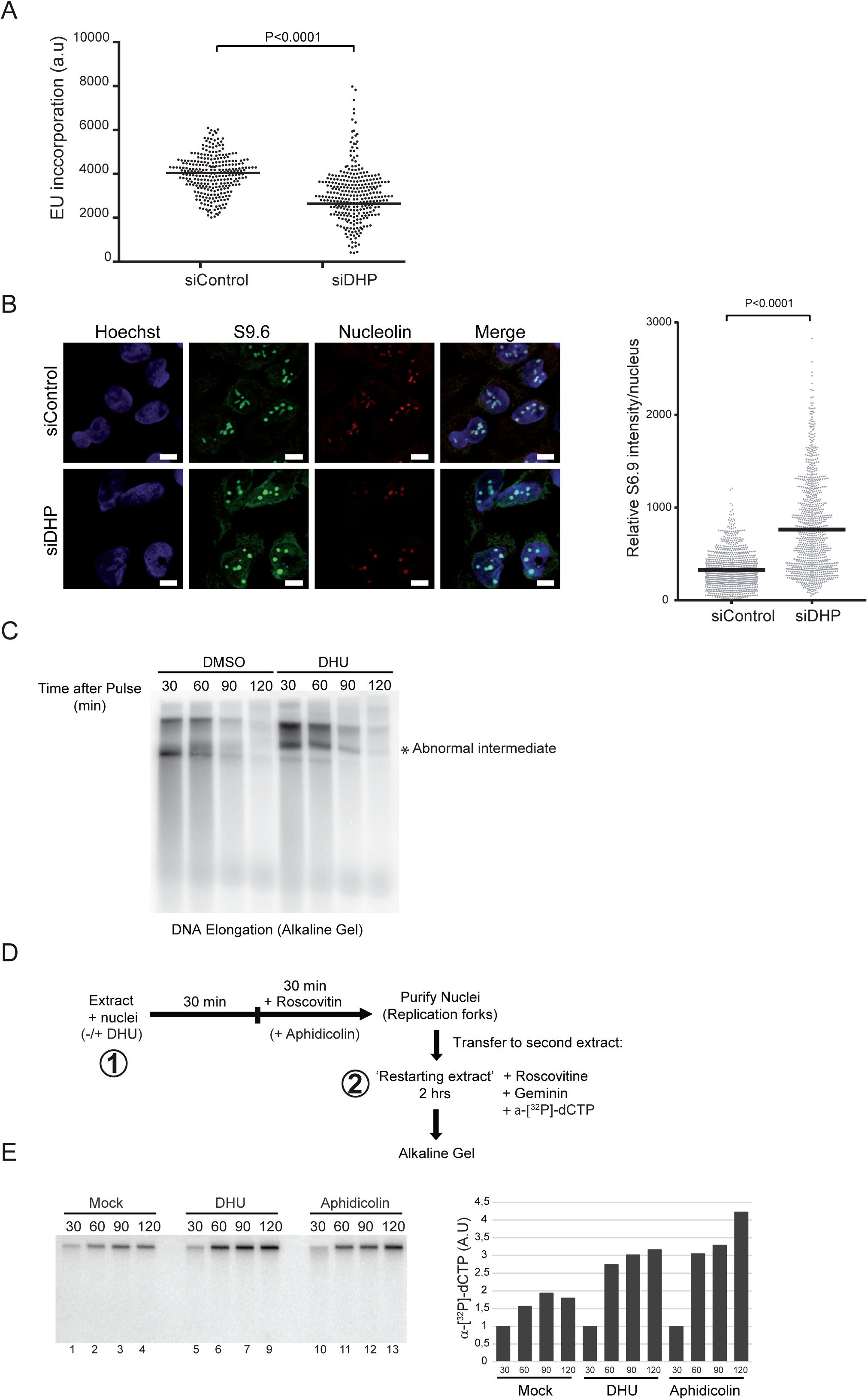
Dihydropyrimidines induce transcriptional stress and yield abnormal DNA replication intermediates. (A) Graphic representation of global transcriptional activity visualized by 5-ethynyl uridine (EU) incorporation. U-2 OS cells transfected with control and anti-DHP siRNA (siDHP-SP) were labelled with EU for 20 min before fixation. The EU intensity of 100 cells from two independent biological replicates was measured by fluorescence microscopy. The bar dissecting the data points represents the median of EU intensity. Differences between distributions were assessed with the Mann-Whitney rank sum test. (B) Immunofluorescence staining with S9.6 and nucleolin antibodies of DHP-depleted or control U-2 OS cells (siDHP-SP). DNA was stained by Hoechst. Bars indicate 10 µm. Right panel: The graph shows the median of S9.6 signal intensity par nucleus after nucleolar signal removal. More than 1000 cells from two independent biological replicates were considered. Differences between distributions were assessed with the Mann-Whitney rank sum test. (C) DNA synthesis reactions (control: DMSO; DHU: 15 mM) were pulse-labeled for 30 min with α-[^32^P]-dCTP at the indicated times during the course of a two-hours reaction. Replication products were purified and resolved by electrophoresis through a 1.2 % agarose gel in denaturing conditions. (*) abnormal replication intermediate. One representative experiment is shown from two biological replicates. (D) Experimental scheme: Sperm nuclei were added to Xenopus egg-extract (in presence or not of 7.5 mM DHU dissolved in water) and incubated at 23 °C to allow origins firing and replication initiation. After 30 min incubation, the firing of new replication origins was blocked with roscovitine (0.5 mM). Replicating nuclei were then isolated after 60 min of incubation and transferred to a second extract (restarting extract) supplemented with roscovitine (0.5 mM) and Geminin (60 mM) to block the firing and the assembly of novel origins, respectively. DNA synthesis reactions were pulse-labeled with α-[^32^P]-dCTP during incubation in the second extract. (E) Replication products were resolved by 1 % alkaline agarose gel electrophoresis and revealed by autoradiography. Lanes 1-4: Mock treated extracts; Lanes 5-9: incubation in the first extract was performed in the presence of 7.5 mM DHU. Lanes 10-13 serve as positive controls: after 30 min incubation in the first extract, DNA synthesis was blocked with aphidicolin (100 ng/µl). Right panel: Histogram representing the quantification of the gel by image J of replication products (arbitrary unit). One representative experiment is shown from two biological replicates (See supplementary figure S3).

### Dihydropyrimidine accumulation induces abnormal DNA replication products independently of transcription

Because interference between transcription and DNA replication is an important endogenous source of DNA replication stress (Tuduri et al., 2009), we wanted to know if dihydropyrimidine metabolites can directly interfere with the process of DNA replication independently of transcription activity. To test this, we used a *cell-free* DNA replication system derived from Xenopus eggs in which transcription is inactive. In this system, a circular single-stranded DNA is converted into a double-stranded DNA via priming and elongation of DNA chains in a semiconservative manner. The replicated DNA is then assembled into chromatin leading to the formation of supercoiled DNA (Mechali and Harland, 1982). DNA replication was measured by the incorporation of the radioactive nucleotide precursor α^32^P-dCTP.

First, we labeled DNA during the course of a two-hour reaction with 30 minutes pulses of α^32^P-dCTP, as indicated, and analyzed DNA replication products by alkaline agarose gel electrophoresis. In these denaturing conditions, irreversibly denatured DNA produced in control extracts was replaced by an abnormal replication intermediate in extracts supplemented with 5, 6-dihydrouracil (Figure 4C). This novel replication product was visible from the earliest stages of the replication reaction (Figure 4C). This observation indicate that dihydropyrimidines interfere directly with the process of DNA chain elongation in Xenopus egg-extracts. Moreover the dNTPs pool is not limiting during DNA replication in xenopus egg extracts. Therefore, in this system, the possibility that DHU may imbalance the pool of dNTP pool is eliminated.

To confirm that 5, 6-dihydrouracil interferes with chromosomal DNA synthesis, we designed a multistep chromatin transfer experiment to confirm that 5, 6-dihydrouracil directly perturbs DNA replication (Figure 4D). First, we performed a standard chromatin DNA replication reaction in Xenopus egg-extracts using demembranated sperm nuclei (Aze et al., 2017; Errico et al., 2007; Trenz et al., 2006). Interference with the progression of replication forks, for example using the replicative DNA polymerase inhibitor aphidicolin, is expected to yield incomplete DNA replication intermediates that can prime DNA synthesis during the course of a second DNA replication reaction. To obtain evidence for the formation of aborted replication intermediates in extracts supplemented with 5, 6 dihydrouracil, we purified and transferred the replicated, or partially replicated, chromatin to a second extract supplemented with both Geminin, to block the licensing of new origins of replication, and the CDK2 inhibitor roscovitine, to block the firing of new origins. In this situation, DNA synthesis is the result of priming of pre-existing replication intermediates. The transfer of nuclei from a replication reaction carried out in the presence of aphidicolin to a second extract unable to fire new origins led to a significant increase in α^32^P-dCTP incorporation in comparison with mock-treated nuclei (Figure 4E and S3A, compare lanes 1-4 with lanes 10-13), indicative of replication fork restart and/or DNA repair activities. Likewise, in comparison with mock-treated nuclei, addition of 5, 6-dihydrouracil during the first replication reaction yielded an increased incorporation of α^32^P-dCTP in the restarting extract (Figure 4E and S3D, lane 5-9), indicating that DNA replication in the presence of 5, 6-dihydrouracil generates DNA intermediates that prime DNA synthesis. Collectively, these data indicate that dihydropyrimidine metabolites directly interfere with the process of DNA replication.

### Dihydromyricetin induces DNA replication stress

We noticed a report suggesting that the plant flavonoid dihydromyricetin is a competitive inhibitor of a putative dihydropyrimidinase from *Pseudomonas aeruginosa* (Huang, 2015) (Figure 1A). Human and *P. aeruginosa* dihydropyrimidinases are predicted to fold into a similar structure (Huang, 2015). Since DHP activity is a good marker of tumorigenicity and a candidate target for cancer therapy (Naguib et al., 1985), we wanted to test if dihydromyricetin also inhibits the human DHP. We expressed and purified recombinant human DHP to near homogeneity (Figure S3B) and assessed its activity by measuring the decomposition of 5, 6-dihydrouracil using high performance liquid chromatography. In an experimental system containing purified dihydropyrimidinase (0.2 μM) and 5,6 dihydrouracil as a substrate (50 μM), dihydromyricetin inhibited dihydropyrimidinase activity with a half maximal inhibitory concentration (IC_50_) value of about 6 μM (Figure 5A). Dihydromyricetin is a versatile molecule (Li et al., 2017). Notwithstanding the pleiotropic actions of dihydromyricetin, we analyzed the phenotypic consequences of exposing U-2 OS cells to 20 μM of dihydromyricetin. This treatment increased the intracellular molar ratio of dihydrouracil/uracil (Figure 5B), induced RPA32 phosphorylation and p53 stabilization (Figure 5C), the formation of RPA32 nuclear foci and (Figure 5D), and the slowing of DNA replication forks (Figue 5E). These data indicate that the treatment of cells with dihydromyricetin induces replication stress phenotypes that are similar to the phenotypes of DHP-knockdown cells.

**Figure 5.**
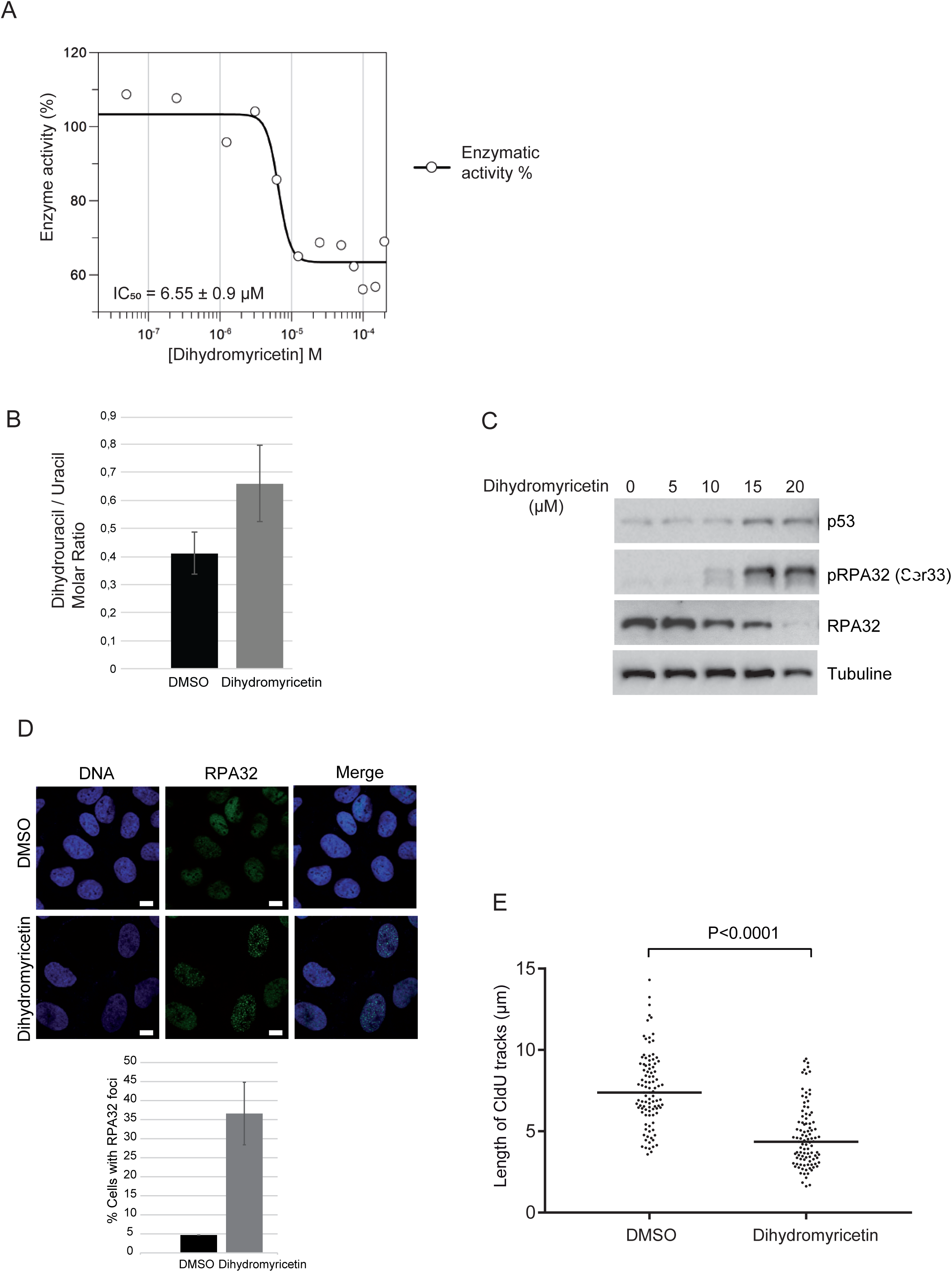
Dihydromyricetin induces DNA replication stress. (A) IC_50_ determination of dihydromyricetin for dihydropyrimidinase (0.2 μM) using dihydrouracil (50 µM) as a substrate. One representative experiment is shown from three biological replicates. (B)Molar ratios of dihydrouracil versus uracil measured in U-2 OS cells treated with 20 µM dihydromyricetin for 16 hr. Data from three independent biological replicates, with three technical replicates for each, are represented as mean +/-S.E.M (C)Western blot analysis with the indicated antibodies of U-2 OS whole-cell extracts treated with dihydromyricetin for 16 hr at the indicated concentrations. One representative experiment is shown from two biological replicates. (D)RPA32 immunofluorescence staining of U-2 OS cells treated with DMSO or 20 µM dihydromyricetin for 16 hr. Bars indicate 10 µm. DNA was stained by Hoechst. Right panel: Histogram representation of the percentage of RPA32 foci-positive cells in a population of 100 cells. Data from three independent biological replicates are represented as mean +/-S.D. (E)Graphic representation of replication track lengths measured in μm (y axis) in control and U-2 OS cells treated with 20 µM of dihydromyricetin for 16 hr. The bar dissecting the data points represents the median of 100 tracts length from one biological replicate. Differences between distributions were assessed with the Mann-Whitney rank sum test.

### Suppression of dihydropyrimidinase activity induces DNA-protein crosslinks

Next, we sought to evaluate whether dihydropyrimidines inhibit DNA-templated processes via the formation of DNA adducts. First, we asked if the accumulation of these metabolites triggers the recruitment of translesion DNA polymerases to chromatin. Suppression of DHP induced chromatin recruitment of the translesion DNA polymerase η that can bypass replication-blocking lesions such as UV photoproducts and oxidized bases (Kannouche et al., 2004; Zlatanou et al., 2011), but not DNA polymerase κ, which has a different specificity for DNA lesions than polymerase η (Figure 6A and 6B). We also observed monoubiquitinated PCNA in the chromatin fraction of DHP-depleted cells (Figure 6A), a posttranslational modification that facilitates the interaction of TLS polymerases with PCNA (Sale et al., 2012). Consistent with this observation, addition of 5, 6-dihydrouracil in Xenopus eggs extracts induced a pronounced accumulation of TLS pol η on chromatin after 2 hours of incubation, when the replication process was completed (Figure 6C). In light of these observations, we propose that the accumulation of dihydropyrimidines or their decomposition products may induce bulky DNA adducts.

**Figure 6.**
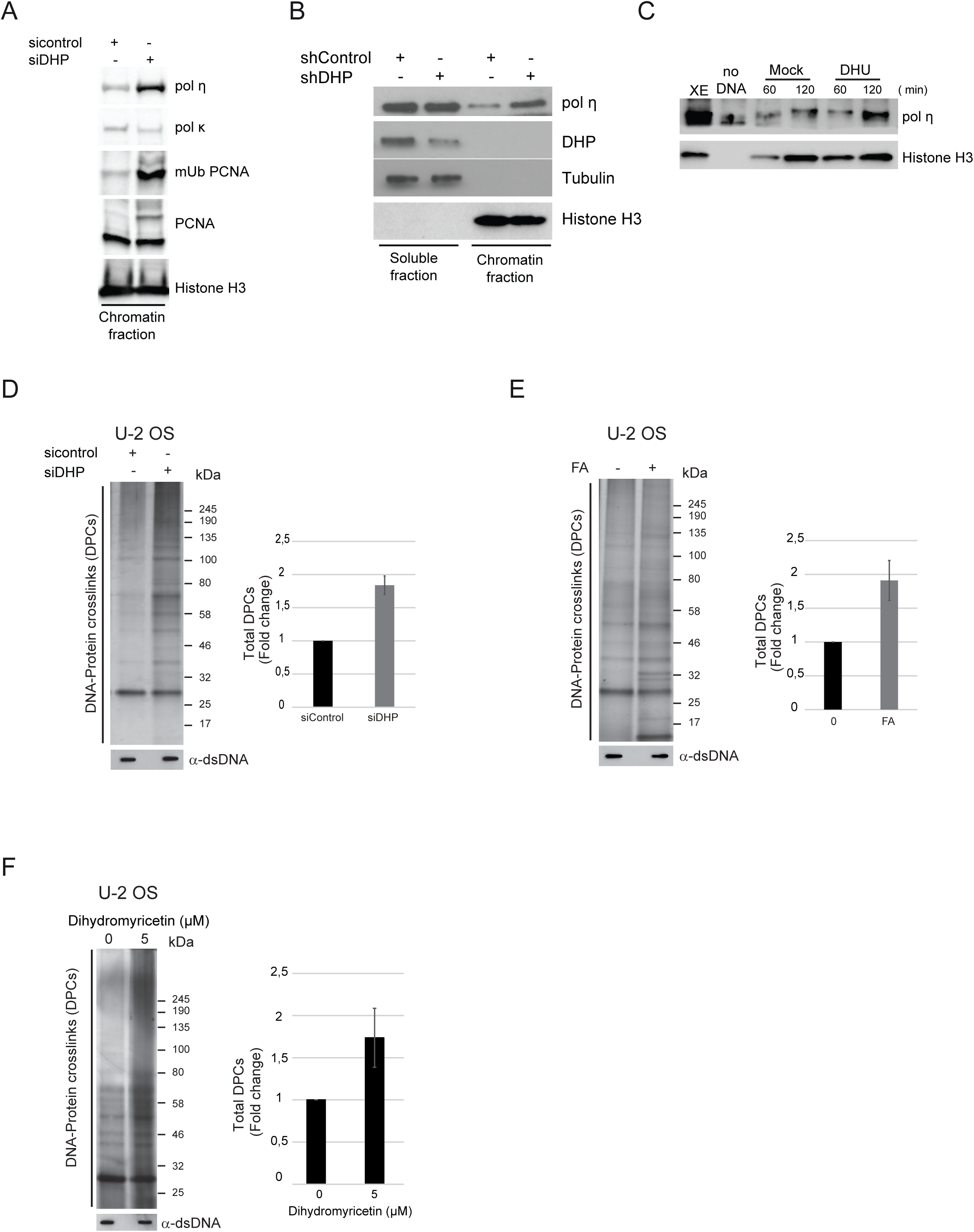
Dihydropyrimidines metabolites induce DNA-protein crosslinks lesions. (A) The chromatin fraction of control and DHP knockdown U-2 OS cells (siDHP) was subjected to western blot analysis with the indicated antibodies. *Histone H3* was used as *loading control.* One representative experiment is shown from two biological replicates. (B) DHP knockdown HEK293T cells were subjected to subcellular fractionation and probed by western blotting with the indicated antibodies. *Tubulin and histone H3* were used as *loading controls* for the cytoplasmic and chromatin *fractions*, respectively. (C) Chromatin extracts from nuclei incubated in control and DHU (7.5 mM) containing extracts for 60 and 120 min were subjected to western blot analysis with the indicated antibodies. *Histone H3* was used as *loading control.* One representative experiment is shown from two biological replicates. (D) Total DPC levels in U-2 OS cells transfected with control or anti-DHP siRNA (siDHP) visualized by silver staining. Right panel: Histogram representing the quantification of DPC levels normalized to total DNA amount by image J. Three independent biological replicates are averaged in the bar graphs. Error bars represent +/-S.D. (E) Total DPC levels in U-2 OS cells treated or not with 1 mM FA for 2 hr visualized by silver staining. Right panel: Histogram representing the quantification of DPC levels normalized to total DNA amount by image J. Three independent biological replicates are averaged in the bar graphs. Error bars represent +/-S.D. (F)Total DPC levels after U-2 OS cells treatment with DMSO or 5 µM of Dihydromyricetin for 16 hr visualized by silver staining. Right panel: Histogram representing the quantification of DPC levels normalized to total DNA amount by image J. Three independent biological replicates are averaged in the bar graphs. Error bars represent +/-S.D.

A variety of endogenous metabolites, environmental and chemotherapeutic DNA damaging agents induce covalent DNA-protein crosslinks (Tretyakova et al., 2015). We use the RADAR assay (rapid approach to DNA-adduct recovery) to test if the suppression of dihydropyrimidinase activity yields DNA-protein crosslinks (DPCs). We isolated genomic DNA and quantified it using Qubit fluorometric quantitation to ensure that DPC analyses were performed using equal amounts of material. Next, we digested DNA with benzonase, resolved DPC by SDS/PAGE and detected them by silver staining. An increase in total DPCs was consistently detected after suppression of DHP using distinct siRNA and shRNA molecules, in U-2 OS and in HEK293T cells (Figure 6D and S4A). The level of DPCs in DHP-depleted cells was comparable to that of U-2 OS cells exposed to formaldehyde (Figure 6E). In addition, treatment of U-2 OS cells with dihydromyricetin increased by 2-folds the amount of total DPCs (Figure 6F). These data suggest that dihydropyrimidines metabolites induce the formation of covalent bonds between proteins and DNA.

A large diversity of proteins can be crosslinked to DNA. As the TLS polymerase η functions at blocked replication forks and accumulates in the chromatin fraction of DHP-depleted cells, we examined if DNA polymerase η was crosslinked to DNA. The level of covalently trapped DNA polymerase η increased in DHP-depleted cells (Figure 7A) and in cells exposed to dihydromyricetin (Figure 7B), suggesting that dihydropyrimidines covalently trap DNA polymerase η on DNA.

**Figure 7.**
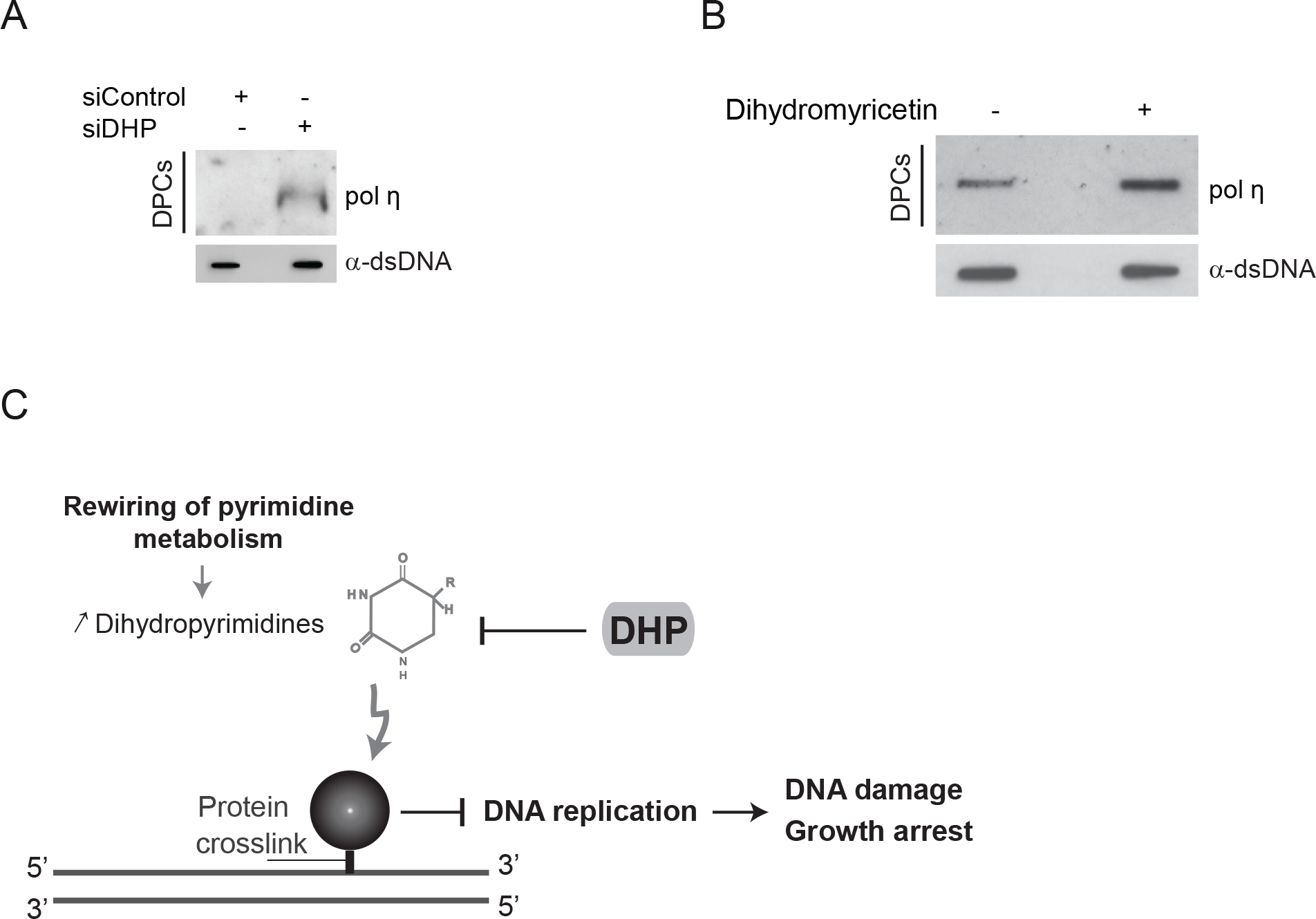
Dihydropyrimidines accumulation induces polymerase η-DNA adducts. (A) Western blot analysis of crosslinked DNA polymerase η in total DPC extracts from U-2 OS cells transfected with control or anti-DHP siRNA (siDHP) and the corresponding DNA quantification. One representative experiment is shown from two biological replicates. (B) Slot-blot showing crosslinked DNA polymerase η in total DPC extracts from U-2 OS cells treated with 20µM dihydromyricetin for 16 hr and the corresponding DNA quantification. (C) Model: The accumulation of Dihydropyrimidines in cancer cells induces DNA replication stress via the formation of DNA-Protein crosslinks (DPCs).

## Discussion

The activity of dihydropyrimidinase is high in the liver and the kidney (van Kuilenburg et al., 2006), absent in the majority of healthy human tissues, and re-expressed in a variety of human carcinomas (Naguib et al., 1985). In this study, we show that dihydropyrimidinase (DHP) is an essential enzyme in transformed cell lines.

Suppression of DHP induced the slowing of DNA replication forks, the accumulation of single-stranded DNA, the activation of ATR signaling, the accumulation of RNA/DNA hybrid structures and the inhibition of transcription. Multiple lines of evidence obtained using orthogonal experimental methods lead us to conclude that dihydropyrimidines are potentially cytotoxic metabolites: (i) Depletion of DHP by RNA interference inhibited DNA replication and transcription; (ii) DNA replication stress in DHP-depleted cells was reversed by suppression of dihydropyrimidine dehydrogenase, the enzyme that produces dihydropyrimidines; (iii) Inhibition of DHP activity with dihydromyricetin phenocopied the defects of DHP-depleted cells; (iv) Addition of dihydropyrimidines in the cell culture medium of transiently permeabilised cells induced markers of DNA replication stress and stabilized p53; (v) Dihydropyrimidines directly altered DNA replication products synthesized in Xenopus egg extracts.

We do not exclude that the observed reduction of dNTPS levels may contribute to the DNA replication stress phenotype of DHP - depleted cells. However, the pool of dNTPs increases in S phase. Thus, the apparent reduction of the pool of dNTPs may simply reflect the reduction of the proportion DHP-depleted cells in S phase. Furthermore, previous studies have shown that the rate of replication fork progression does not correlate directly with the global concentration of dNTPs (Kumar et al., 2010; Techer et al., 2016), as the latter does not reflect the amount of nucleotides available to the replication machinery. By contrast, the length of replication tracks is determined directly by the accumulation of DNA lesions in the template DNA and by p53 activation (Techer et al., 2016). Several lines of evidence suggest that it is unlikely that alterations of the pool of dNTPs determine the phenotypes of DHP-depleted cells. First, the length of replication tracks in DHP-depleted cells remain shorter than in control cells after saturation of the cell culture medium with exogenous nucleotides. Second, the pool of dNTPs is not limiting in Xenopus egg extracts, yet supplementation of Xenopus egg extracts with dihydropyrimidines interfered with DNA replication. Third, transcription does not depend on dNTPs precursors, yet, the accumulation of dihydropyrimidines also inhibited RNA synthesis. The accumulation dihydropyrimidines induced p53 stabilization and DNA damage, two parameters that determine directly the length of replication tracks (Techer et al., 2016).

Cellular metabolites and environmental agents generate a range of structurally diverse protein-DNA crosslink that precipitate the loss of cellular functions, including transcription and DNA replication (Tretyakova et al., 2015). The data presented here reveal that dihydropyrimidines induce the formation of DNA-protein crosslinks, including covalently trapped DNA polymerase η. Yet, we did not elucidate how dihydropyrimidines yield protein-DNA adducts. Some evidence suggests that hydrouracil and its derivatives could be incorporated into ribonucleic acids (Mokrasch and Grisolia, 1958, 1960), but it is not clear whether salvage pathways can convert 5,6-dihydrouracil and 5,6-dihydrothymine into nucleosides or deoxynucleosides. During the course of this study, we did not detect any direct evidence of dihydropyrimidines incorporation into DNA. Dihyropyrimidines are non-coding bases that have lost their planar structure as a consequence of the saturation of the C5-C6 double bound (Lindahl, 1993). Above physiological pH, and more slowly at physiological pH, these saturated bases can further decompose into fragments of bases (Lin et al., 2014). Some decomposition products could be genotoxic. Alternatively, metabolites alterations by chemical side reactions are widespread (Lerma-Ortiz et al., 2016). Dihydropyrimidines may react with oxidants or other metabolites to form potent DNA damaging agents (Wang et al., 2013).

A deficiency in DHP activity yields clinical symptoms that are consistent with dihydropyrimidines exerting toxic effects. Individuals carrying bi-allelic mutations in DHP accumulate high levels of 5, 6-dihydrouracil and 5, 6-dihydrothymine in urine, blood, and cerebrospinal fluids. DHP deficiency can remain asymptomatic, but most patients present neurological abnormalities including mental retardation, hypotonia and seizures (Putman et al., 1997; Sumi et al., 1996; van Kuilenburg et al., 2010; van Kuilenburg et al., 2007). DHP deficiency also manifests with growth retardation, dysmorphic features and gastrointestinal abnormalities (Assmann et al., 1997; Hamajima et al., 1998; Henderson et al., 1993).

In the context of carcinogenesis, the rewiring of the pyrimidine degradation pathway appears as a metabolic adaptation that supports tumor progression. Most cultured cells require glutamine for TCA cycle anaplerosis that yields precursors for several biosynthetic pathways, including nucleotides, which are necessary for tumor growth (Lunt et al., 2015; Vander Heiden and DeBerardinis, 2017). The accumulation of the pyrimidine degradation products dihydropyrimidines facilitate directly the epithelial mesenchymal transition (EMT) (Shaul et al., 2014). The role of dihydropyrimidines in cancer progression towards metastasis could be linked with the genotoxic potential of these metabolites. Endogenous DNA damage and DNA replication stress induce genomic instability and thereby accelerate the acquisition of growth promoting properties. Furthermore, key sensors and mediators of the DNA damage response regulate the EMT-associated transcription factor ZEB1 (Liu et al., 2015; Park et al., 2016; Zhang et al., 2014a). An overload of DNA lesions, however, will trigger cell death. The previously unrecognized cytotoxicity of dihydropyrimidines described here implies that a tight equilibrium between pyrimidines synthetic and pyrimidine degradation activities is required for the proliferation of some cancer cells. We propose that DHP fulfills the function of a sanitization enzyme required for epithelial cancer cells to mitigate the toxicity of dihydropyrimidines (Figure 7C).

Dihydropyrimidinase is a potential target for cancer chemotherapy (Naguib et al., 1985). We report that dihydromyricetin inhibits the activity of recombinant human DHP. Intriguingly, U-2 OS cells exposed to dihydromyricetin exhibited cellular phenotypes similar to that of DHP-depleted cells: (i) accumulation of protein-DNA crosslinks; (ii) replication forks slowing; and (iii) induction of markers of DNA replication stress. The later observation confirms a previous study showing that in hepatocellular carcinoma, dihydromyricetin elicits p53 stabilization and activates the G2/M checkpoint via ATM/ATR signaling (Huang et al., 2013). Dihydromyricetin is a versatile flavonoid from the Chinese pharmacopeia. It scavenges reactive oxygen species, probably interacts with many molecules and proteins and has a variety of biological activities (Li et al., 2017). Dihydromyricetin induces cell cycle arrest and apoptosis in human gastric cancer cells, hepatocellular carcinoma and melanoma cells, without cytotoxicity to normal cells (Huang et al., 2013; Ji et al., 2015; Zeng et al., 2014; Zhang et al., 2014b). It inhibits the growth of prostate cancer in mice (Ni et al., 2012). This study shows that DHP activity is a cellular target of dihydromyricetin. It offers a conceptual framework for further exploring the potential therapeutic utility of targeted inhibition of DHP.

## Acknowledgment

We thank Helena Sapede, Touffic Kassouf for technical help, Ketan J Patel, Bruno Vaz and Kristijan Ramadan for helpful discussion, and members of the laboratory for critical reading of the manuscript. We acknowledge the imaging facility MRI, member of the national infrastructure France-BioImaging infrastructure supported by the French National Research Agency (ANR-10-INBS-04, «Investments for the future»), and the SIRIC Montpellier Cancer (grant INCa-DGOS-Inserm grant 6045). This work was supported by the Fondation ARC pour la recherche sur le cancer (grant 060509 to A.C.) and by grants from l’Institut National du Cancer (grant INCa121770 to J.M. and A.C.), MSD Avenir (to A.C.), USA NIH (grants AI049781 and GM104198 to B.K.), and INSERM Plan Cancer (to D.M.).

## Methods

### Cell lines, plasmids and chemicals

U-2 OS, HEK293T and MCF7 were grown under standard conditions in Dulbecco’s modified Eagle’s medium (DMEM) (Invitrogen) supplemented with 10 % fetal bovine serum (FBS) and 1 % *penicillin*/*streptomycin* (P/S). HCT116 (Horizon Discovery Ltd.) were cultured in McCoy’s 5A modified Medium (Sigma-Aldrich) supplemented with 10 % FBS and 1 % P/S. XG1 and XG19 IL6 dependent human myeloma cell lines (HMCLs) were obtained as previously described (Moreaux et al, 2011). AMO-1 and OPM2 were purchased from DSMZ (Braunsweig, Germany). These HMCLs were routinely maintained in RPMI 1640 and 10% fetal calf serum (FCS; Biowittaker, Walkersville, MD), supplemented with 3 ng/mL IL-6 (Peprotech, Rocky Hill, NJ) for IL6 dependent cell lines. HMCLs were authenticated according to their short tandem repeat profiling and their gene expression profiling using Affymetrix U133 plus 2.0 microarrays deposited in the ArrayExpress public database under accession numbers E-TABM-937 and E-TABM-1088. Dihydrouracil, Uracil, Dihydromyricetin, Aphidicolin and Roscovitine were purchased from Sigma-Aldrich. Formaldehyde was purchased from VWR chemicals. Embryomax nucleosides (100X) (cytidine, 0.73 g/liter; guanosine, 0.85 g/liter; uridine, 0.73 g/liter; adenosine, 0.8 g/liter; thymidine, 0.24 g/liter) was purchased from Millipore. pDONR223-DPYS was obtained through MGC Montpellier Genetic Collections and cloned into destination vectors using gateway technology (Invitrogen).

### Antibodies

Primary antibodies were purchased from Abcam (Histone H3, p53, DNA polymerase eta, nucleolin), Bethyl Laboratories (RPA32-Ser33, RPA32-Ser4/S8, DNA polymerase kappa), Calbiochem (RPA32), Cell Signaling Technology (Chk1-Ser345, Ubiquityl-PCNA (Lys164)), ProteinTech Group (DPYS), Santa Cruz Biotechnology (Chk1, HA), Sigma-Aldrich (α-Tubulin, DPYD, PCNA). Xenopus polymerase η antibody was previously described (Kermi et al., 2015). The Mouse anti-RNA-DNA hybrid S9.6 hybridoma was purchased from ATCC. Orc2 antibodies were kindly provided by Dr. Marcel Mechali (Institute of Human Genetics). Secondary antibodies (anti-rabbit-HRP and anti-mouse-HRP) were from Promega

### DHP bacterial protein expression and purification

DPYS was cloned into the pET-28a (+) (Novagen) vector containing an N-terminal 6xHis tag. The protein was overexpressed in E.coli BL21(DE3) host cells and induced by 1 mM Isopropyl β-D-1-thiogalactopyranoside (IPTG) (Sigma-Aldrich) for 3 hr in presence of 1 mM ZnCl2. Cells were lysed with Buffer A (50 mM Potassium Phosphate pH 7.5, 150 mM NaCl, 0.1 % NP40, 15 mM Imidazol (Sigma-Aldrich)) and protease inhibitors (Roche). Extracts were incubated for 30 min at 4 °C and harvested at 28,000 rpm for 1 hr. The soluble supernatant fraction was purified on a 5 ml HisTrap HP column (GE Healthcare) using the AKTA protein purification system (GE Healthcare). The column was washed with ten column volumes of Buffer A with 60 mM Imidazol. Bound protein was eluted from the column using Buffer A with 250 mM Imidazol. The fractions corresponding to each peak in the chromatogram were dialysed against buffer containing 50 mM Tris HCl pH 7.5, 150 mM NaCl, 1 mM DTT and 10 % glycerol.

### RNA interference and transfection

ON-TARGET plus siRNA Human DPYS siRNA SMARTpool (siDHP-SP) (L-008455-00), ON-TARGET plus siRNA Human DPYS (siDHP) (J-008455-07) (GCACAGAUGGCACUCACUA), siGENOME SMARTpool Human DPYD (M-008376-02) and ON-TARGET plus Non-Targeting Pool (D-001810-10) were purchased from Dharmacon. siDHP-2 (GAAUAGCUGUAGGAUCAGATT) was purchased from Eurofins MWG. DPYS MISSION shRNA plasmid (TRC0000046747) (TGTGGCAGTTACCAGCACAAA) and (TRC0000046744) (CTAATGATGATCTAACCACAA) was from Sigma-Aldrich. siRNA and shRNA transfections were performed using INTERFERin (Polyplus) and Lipofectamine 2000 (Invitrogen) reagents, respectively and the different cDNA plasmids were transfected using jetPEI or jetPRIME reagent (Polyplus).

### Small-Scale Chromatin Fractionation Assay and western blotting

As described (Wysocka et al, 2001), cells were collected, washed with phosphate-buffered saline (PBS), and resuspended in buffer A (10 mM HEPES [pH 7.9], 10 mM KCl, 1.5 mM MgCl_2_, 0.34 M sucrose, 10 % glycerol, 1 mM dithiothreitol, and protease inhibitors (Roche)). Triton X-100 was added (0.1 % final concentration), the cells were incubated on ice for 5 min, and nuclei were collected by centrifugation (5 min, 1,300 × g, 4 °C). The supernatant (Cytosolic fraction) was clarified by high-speed centrifugation (5 min, 20,000 × g, 4 °C), and the supernatant (Cytosolic fraction) was collected. The nuclei were then washed once in buffer A and lysed for 30 min in buffer B (3 mM EDTA, 0.2 mM EGTA, 1 mM dithiothreitol, and protease inhibitor (Roche)), and insoluble chromatin and soluble fractions (Nucleosolic fraction) were separated by centrifugation (5 min, 1,7000 × g, 4 °C). The insoluble chromatin fraction was washed twice with buffer B and resuspended in sodium dodecyl sulphate (SDS)-Laemmli buffer and boiled for 10 min. Western blotting was performed using the ECL procedure according to the manufacturer’s instructions (Amersham Bioscience, Inc) using anti-mouse or rabbit-HRP secondary antibodies (Promega)

### Immunofluorescence Staining and ssDNA detection

Cells grown on coverslips were fixed with 3.7 % paraformaldehyde (PFA) in PBS for 15 min at RT followed by a 0.5 % Triton X-100-PBS permeabilization step for 10 min. Cells were then blocked in PBS containing 3 % BSA for 30 min and incubated in the primary antibody and then in the appropriate secondary antibodies Alexa Fluor 488 or Alexa Fluor 555 (Invitrogen), diluted in blocking solution for 1 hr in a humidified chamber at RT. DNA was stained with Hoechst (Invitrogen) and coverslips were mounted on glass slides with Prolong (Sigma-Aldrich).

For ssDNA detection, cells were grown on microscopic slides in 20 µM BrdU for 24 hr. Primary mouse antibody against BrdU in ssDNA was used (BD). All the Microscopic analysis was performed using Zeiss Z2 Axioimager with ApoTome. ImageJ was used for picture processing and quantification of S9.6 mean intensity.

### DNA Fiber Labeling

DNA fiber spreads were prepared as described previously (Jackson & Pombo, 1998). Cells were labelled with 25 µM IdU (5-*iodo*-2′-deoxyuridine), washed with warm media and then exposed to 50 µM CldU (5-*Chloro*-2′-deoxyuridine). Cells were lysed with the spreading buffer (200 mM Tris-HCl pH 7.5, 50 mM EDTA and 0.5 % SDS) and DNA fiber were stretched onto glass slides. The DNA fibers were denatured with 2.5 M HCl for 1 hr, washed with PBS and blocked with 2 % BSA in PBS-Tween 20 for 60 min. IdU replication tracts were revealed with a mouse anti-BrdU/IdU antibody (BD Bioscience) and CldU tracts with a rat anti-BrdU/CldU antibody (Abcam). DNA fibers were uniformly labeled with a mouse anti-human single-stranded DNA antibody (Millipore). The secondary antibodies used for the assay were: alexa fluor 488 anti-mouse antibody (Life technologies), alexa fluor 647 anti-mouse antibody (Life technologies), and Cy3 anti-rat antibody (Jackson Immunoresearch). Replication tracts were analyzed with ImageJ software. The probability that two data sets stem from the same distribution was assayed by a non-parametrical Mann-Whitney test (Prism Software).

### Fluorescence-activated cell sorting (FACS)

Cells were pulse labeled with 10 µM BrdU for 15 min before fixation with ice-cold 100 % ethanol. Then cells were incubated with PBS and 50 µg/ml of RNase A for 1 hr at 37 °C. After treatment with 2N HCl for 30 min, cells were incubated with an anti-BrdU antibody (BD) for 1 hr at RT and then with an FITC-conjugated anti-mouse IgG (Life Technologies) at RT for 30 min. Cells were stained with 25 µg/ml of propidium iodide in PBS and analyzed using a FACSCalibur machine (BD).

### Enzyme assay

5, 6-Dihydrouracil (DHU) was used as the substrate in the standard assay of dihydropyrimidinase. Briefly, 2.5 µg of purified His-tagged DHP was added to 200 µl of 5, 6-dihydrouracil (50 µM) solution containing 50 mM Tris, 50 µM DTT, pH 8.0 in presence of several concentrations of dihydromyricetin, the samples were then incubated at 37°C for 1 hr. An aliquot (100 µl) from each point was taken before incubation as a control without enzymatic reaction. DHU decomposition was monitored by HPLC using a Waters Alliance system connected to a C18 reversed phase Symmetry column (4.6 × 150 mm, 5 µm, Waters). Elution of DHU was achieved by applying an isocratic flow of H_2_O/TFA 0.1% as mobile phase for 15 min using a flow rate of 1 mL/min. For each sample, the column was first washed with 20% CH_3_CN / TFA 0.1% to remove any residual dihydromyricetin and equilibrated again with elution phase for 10 min. Under these conditions DHU was eluted at 2.8 min and detected by absorbance at 230 nm. Quantification of DHU was performed by integration of the corresponding HPLC peak using Empower Pro software. IC_50_ calculation was performed using Grafit-7 (Erithacus Software).

### Intracellular dNTP measurement

For dNTP analysis and quantification, siRNA or shRNA transfected cells were harvested and lysed in iced cold 65 % methanol, and vigorously vortexed for 2 min. Extracts were incubated at 95 °C for 3 min. Supernatants were collected and dried in a speed vacuum. Samples were processed in Kim Baek laboratory for the single nucleotide incorporation assay as described (Diamond et al, 2004).

### Metabolite extraction

Cells pellet (1 million cells) were extracted on dry ice in 0.5 ml cold 70% methanol. The cell mixtures were shaken vigorously on a Vortex mixer for 10 min. These extracts were then centrifuged at 20 000 g at +4°C for 10 min, and the supernatants were transferred into polypropylene tubes for evaporation with a turbovap evaporator (Biotage, France). Dried extracts were reconstituted in 50 µL of mobiles phases A/B; 9/1. Five µL of this sample was injected in the LC-MS/MS system. For LC Analysis: An UPLC Acquity I Class (Waters, France) was used for this study. The chromatographic separation was perform onto an Acquity UPLC BEH HSS T3 column (150 × 2.1mm, 1.8 µm) using a gradient from 0.5%formic acid in water/0.5% formic acid in acetonitrile; 9/1; v/v as initial conditions to 6/4; v/v from 0.5 min to 3 min at a flow rate of 0.3mL/min. The run time was 5 min allowing the system to reach 100% of 0.5% formic acid in acetonitrile to rinse the column and return to initial mobile phase conditions. The autosampler and the column compartment were held at 4°C and 30°C, respectively. Under these conditions, uracil and dihydrouracil displayed a mean retention time of 1.33 and 1.34 min, respectively. For MS Analysis, the UPLC system was coupled to a Waters XEVO™ TQ-XS mass spectrometer (Waters, France) operating in positive ion mode. For pyrimidine detection, the capillary voltage was set to 2.5 kV. The source and desolvation temperatures were held at 150°C and 600 °C, respectively. The cone and desolvation gas flow were set at 150 and 800 L/hr, respectively. The MS data acquisition was performed in multiple reaction monitoring (MRM) mode. Monitored MRM transitions were m/z 112.93>69.96 and 114.98>72.91 for uracil and dihydrouracil, respectively. Range of calibration curves were 0.28-108 and 0.27-110 nmol/cell pellet for uracil and dihydrouracil, respectively.

### Cell viability and colony forming assay

The effect of siRNA on cell proliferation was measured using the CellTiter-Glo^®^ Luminescent Cell Viability Assay Kit (Promega) according to the manufacturer’s protocol or using the MTT cell growth assay. Briefly, siRNA transfected cells were seeded in 96-well plate and four days later, 100 μl CellTiter-Glo^®^ reagent was added to each well that contained 100 μl cell culture medium. Cells were then lysed by shaking in an orbital shaker for 2 minutes, followed by incubation at room temperature for 10 minutes to stabilize the luminescent signal. The luminescent intensity was recorded on a Tristar LB 941 Multimode Microplate Reader (Berthold Technologies). For the MTT assay, siRNA-transfected cells were seeded into 24-well plates and cell growth was documented every 24 hr via a colorimetric assay using a 3-(4,5-dimethylthiazol-2-yl)-2,5-diphenyltetrazolium bromide (MTT) assay (Sigma-Aldrich). Absorbance values were collected at 600 nm using a BioPhotometer (Eppendorf). In each individual experiment, proliferation was determined in triplicate, and the overall experiment was repeated three times. For colony formation analysis, cells were seeded in six-well plates at a density of 5000 cells per well. The medium was changed every 3 days for 10 days until visible colonies formed. Colonies were fixed in methanol for 10 min and stained with crystal violet.

### Transcription assay

siRNA transfected cells were grown on cover slips and incubated with EU for 20 min. Cells were fixed with 4 % paraformaldehyde for 15 min and permeabilized for 20 min in 0.1 % Triton X-100 in PBS. EU incorporation were detected by staining with the Click-it Edu Alexa Fluor 555 azide Imaging Kit (Invitrogen) according to the manufacturer’s instructions and DNA was stained with Hoechst (Invitrogen). The intensity of staining within individual nuclei was quantified using Image J software.

### *X. laevis* egg extracts preparation and DNA replication kinetics

Low Speed Egg extracts (LSE) were prepared as previously described (Lutzmann & Mechali, 2008). M13 replication kinetics was assessed using 500 ng of M13mp18 Single-stranded DNA (New England BioLabs) per 50 µl of LSE supplemented with cyclohexymide (250 µg/ml), an energy regeneration system (1 mM ATP, 2mM MgCl_2_, 10 mM creatine kinase, 10 mM creatine phosphate) and α-[^32^P]-dCTP (0.37 MBq). Chromosomal DNA replication was assessed by adding 1000 demembranated *X.laevis* sperm nuclei per microliters of extract. The mixtures were incubated at 23 °C for the indicated time, then samples were neutralized in 10 mM EDTA, 0.5 % SDS, 200 µg/ml Proteinase K (Sigma-Aldrich) and incubated at 52 °C for 1 hr. Incorporation of radiolabeled deoxynucleotides in DNA was monitored using a Phosphor Imager Typhoon TriO+ (Amersham Biosciences) following agarose gel electrophoresis or alkaline agarose gel electrophoresis of purified DNA.

Sperm chromatin purification was performed as previously described (Recolin B, 2012 NAR). Briefly, egg extracts supplemented with demembranated sperm nuclei were diluted 10-fold in ice cold XB (10 mM Hepes-KOH pH 7.7, 100 mM KCl, 2 mM MgCl_2_, 50 mM sucrose and protease inhibitor) and pelleted at 1500g for 5minutes. Nuclei were washed once in XB buffer and then detergent extracted with 0.1 % NP40 for 5 minutes on ice. Chromatin was recovered after centrifugation and ressuspended in Laemmli buffer for western blot analysis. For the chromatin transfer experiments, chromatin samples were incubated for 30 min in the first extract with the indicated drugs (Figure 5D). Purification for chromatin transfers and isolation of nuclei were performed following the protocol detailed in (Gillespie et al., 2012). Isolated nuclei integrity was verified by microscopy and then transferred into fresh extract supplemented with geminin and Roscovitine (to inhibit new replication events) together with α-[^32^P]-dCTP. After 2 hours, dCTP incorporation was monitored autoradiography on neutral gel as described previously.

### DNA-protein crosslinks isolation and detection

DNA-protein crosslinks isolation DPCs were prepared as described in (Vaz et al., 2016). In brief, 1.5 to 2 × 10^6^ cells were lysed in 1 ml of M buffer (MB), containing 6 M GTC, 10 mM Tris–HCl (pH 6.8), 20 mM EDTA, 4 % Triton X-100, 1 % Sarkosyl and 1 % dithiothreitol. DNA was precipitated by adding 1 ml of 100 % ethanol and was washed three times in wash buffer (20 mM Tris–HCl pH 6.8, 150 mM NaCl and 50 % ethanol) and DNA was solubilized in 1 ml of 8 mM NaOH. A small aliquot of the recovered DNA was digested with 50 µg/ml proteinase K (Invitrogen) for 3 hr at 50 °C and quantified using Qubit dsDNA HS Assay Kit (Invitrogen) according to manufacturer instructions. DNA concentration was further confirmed by slot-blot where the proteinase K digested samples were diluted in TBS buffer and applied to nylon membrane (Hybond N+) followed by immunodetection with antibody against dsDNA. The remaining solubilized DNA was digested with Benzonase (Sigma-Aldrich) for 30 min at 37 °C. Proteins were precipitated by standard Trichloroacetic Acid (TCA) protocol. Finally, the crosslinked-proteins were resuspended with the appropriate buffer and total DPCs were analyzed by Silver Staining (Invitrogen) as recommended by the manufacturer after electrophoretic separation on polyacrylamide gels and specific crosslinked-proteins were immunodetected using Western blot assay. Signals were quantified using Image J software.

### Statistical analysis

The biological replicates are experiments that have been performed independently of each other and not simultaneously (which indicate biological variation). The technical replicates are replicates performed during one biological replicate (which indicate variation of the measuring equipment and protocols).

---

**Figure S1.**
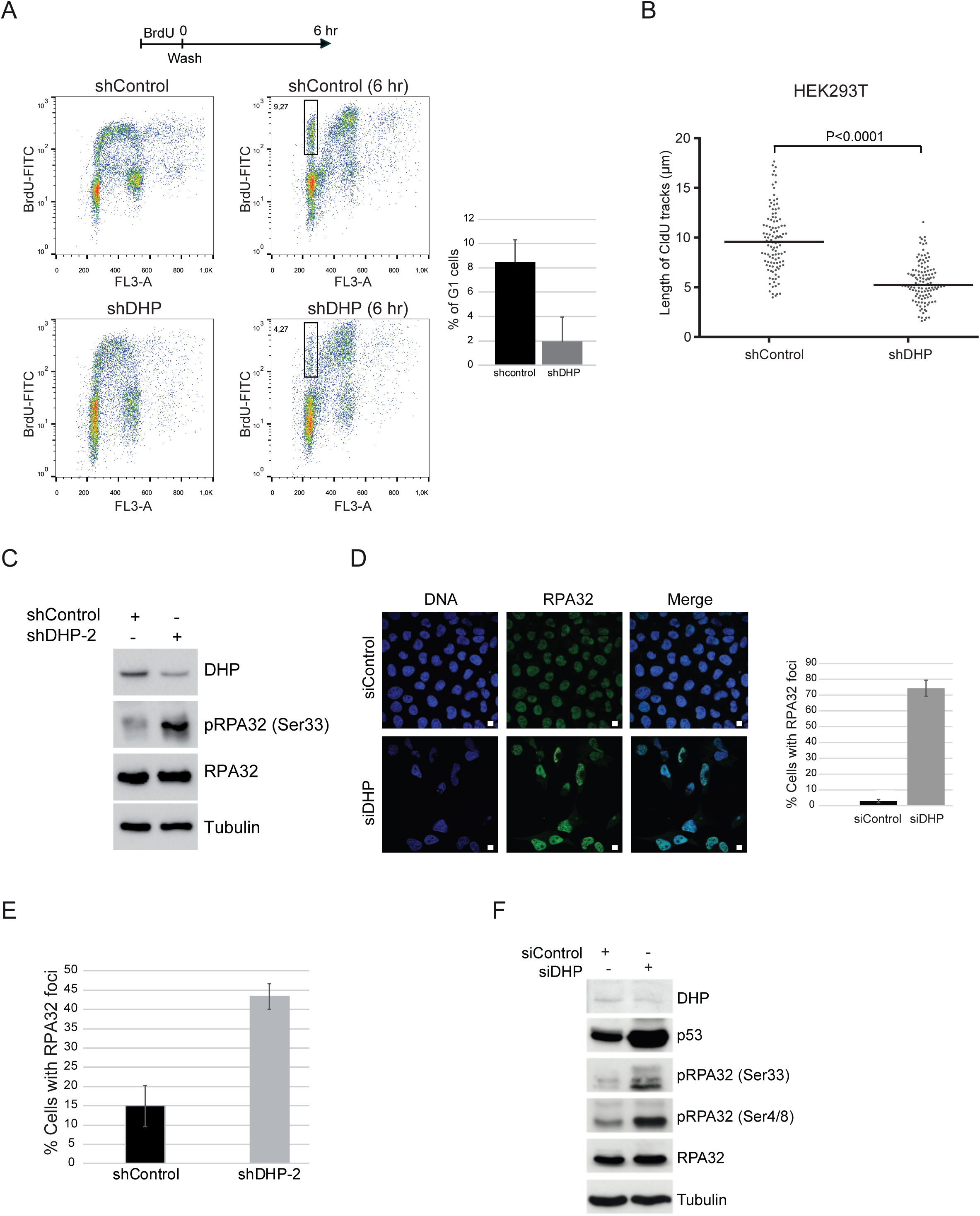
(A) Upper panel: Experimental scheme. HEK293T cells transfected with control or anti-DHP shRNA were pulse-labeled with 20 μM BrdU for 30 min, washed (W) and analyzed by two-dimensional (BrdU/DNA) flow cytometric analysis at the indicated time. One representative experiment is shown from three biological replicates. (B) Graphic representation of replication track lengths in control and HEK293T cells for DHP using a shRNA molecule with a distinct target sequence in DHP. The bar dissecting the data points represents the median of 100 tracts length from one biological replicate. Differences between distributions were assessed with the Mann-Whitney rank sum test. (C) Whole-cell extracts from control and DHP knockdown HEK293T cells (shDHP-2) were analyzed by western blotting with the indicated antibodies. One representative experiment is shown from two biological replicates. (D) RPA32 immunofluorescence staining of control and DHP knockdown U-2 OS cells (siDHP). DNA was stained by Hoechst. Bars indicate 10 µm. Right panel: Quantification of the percentage of RPA32 foci positive cells in a population of 100 cells. Data from three independent biological replicates are represented as mean +/-S.D. (E) Quantification of the percentage of RPA32 foci positive cells in a population of 100 cells of DHP-depleted HEK293T cells (shDHP-2). Data from three independent biological replicates are represented as mean +/-S.D. (F) Whole-cell extracts from control and DHP knockdown U-2 OS cells (siDHP) were analyzed by western blotting with the indicated antibodies. One representative experiment is shown from more than three biological replicates

**Figure S2.**
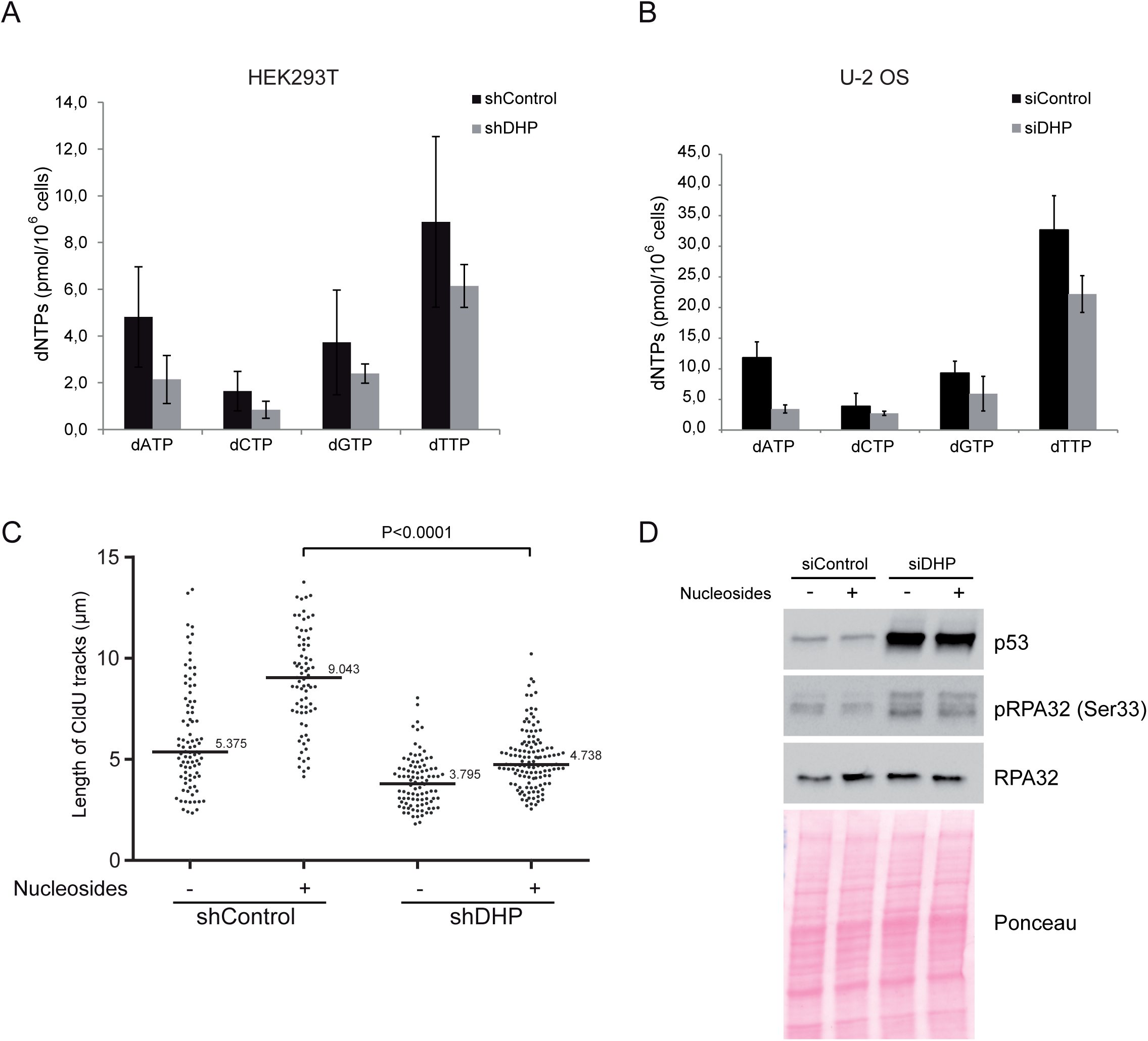
(A)Analysis of dNTP concentrations in control and DHP knockdown HEK293T cells. Data from three independent biological replicates, with three technical replicates for each, are represented as mean +/-S.D. Cellular dNTPs were measured by HIV-RT based dNTP assay (Diamond TL et al, 2004). (B) Analysis of dNTP concentrations in control and DHP knockdown U-2 OS cells. Data from three independent biological replicates, with three technical replicates for each, are represented as mean +/-S.D. (C) Control and DHP knockdown HEK293T cells were supplemented with nucleosides and incubated for 18 hours before DNA fiber analysis of the length of CldU labeled replication tracks, in μm (y axis). The bar dissecting the data points represents the median of 100 tracts length from one biological replicate. (D) Control and DHP knockdown U-2 OS cells (siDHP) were supplemented with nucleosides and incubated for 18 hours before western blot analysis with the indicated antibodies. Ponceau staining was used as loading control.

**Figure S3.**
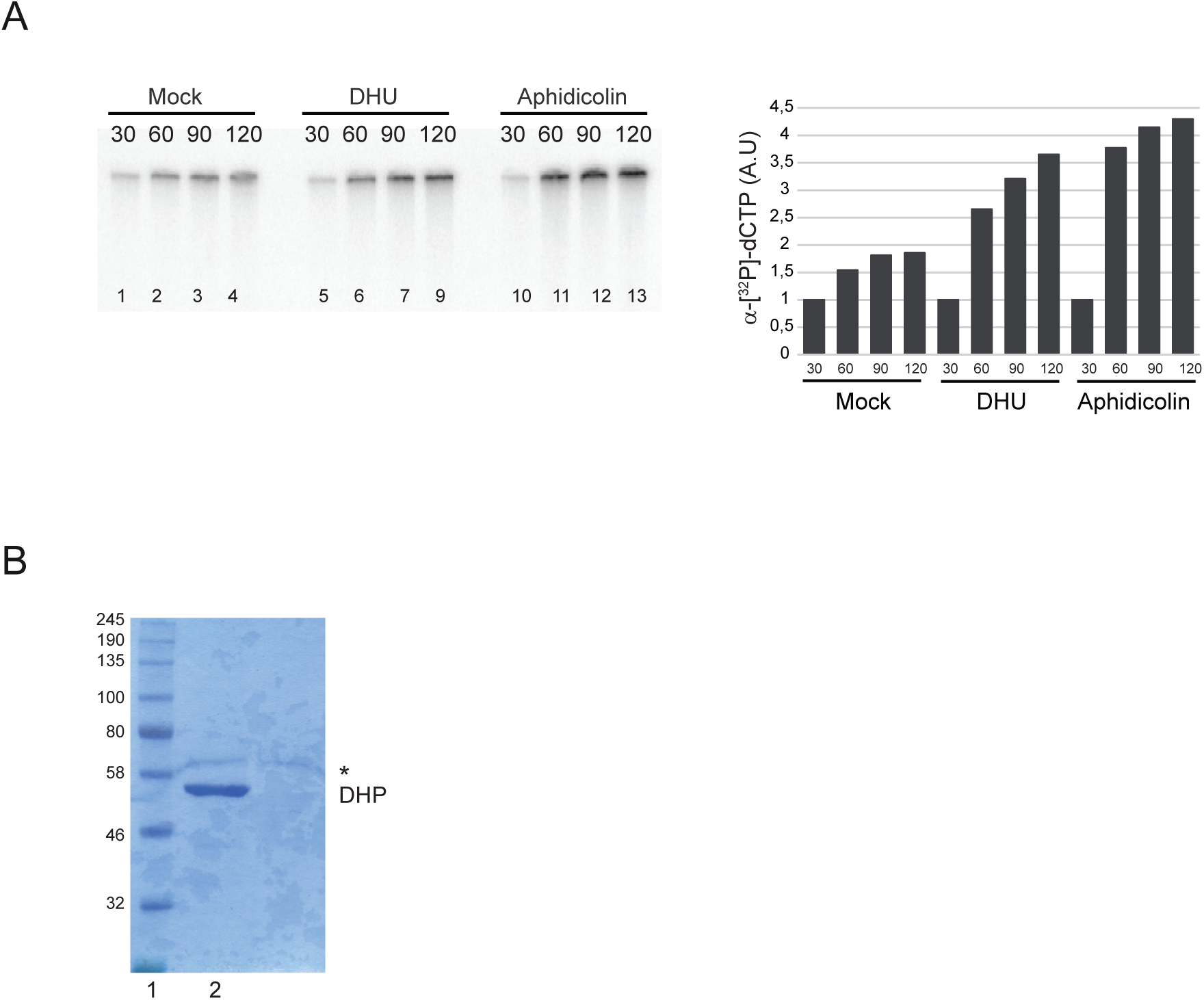
(A) Replicate of chromatin transfer as described in Figure 5E. Replication products were resolved by 1 % alkaline agarose gel electrophoresis and revealed by autoradiography. Lanes 1-4: Mock treated extracts; Lanes 5-9: incubation in the first extract was performed in the presence of 7.5 mM DHU. Lanes 10-13 serve as positive controls: after 30 min incubation in the first extract, DNA synthesis was blocked with aphidicolin (100 ng/µl). Right panel: Histogram representing the quantification of the gel by image J of replication products (arbitrary unit). (B) Purified His-tagged DHP was resolved by SDS/PAGE and stained with Coomassie (lane 2). Size marker (lane 1). * non-specific band.

**Figure S4.**
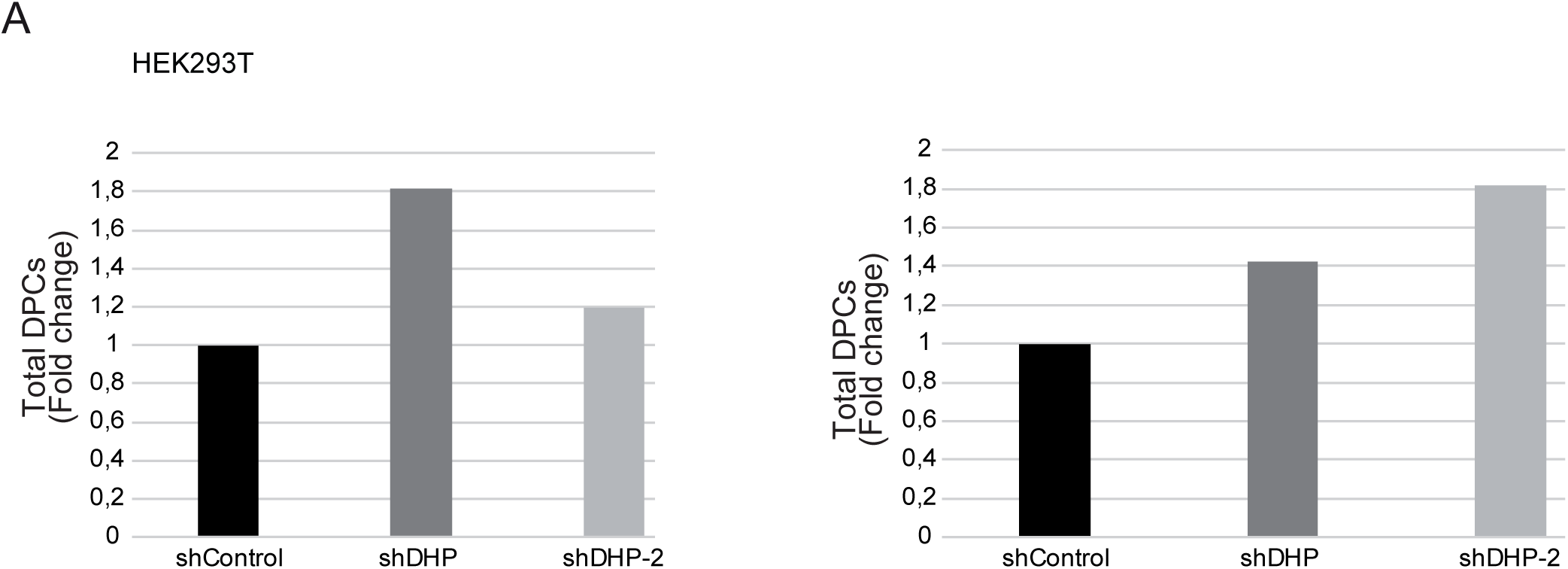
(A) Histogram representing the quantification of DPC levels of HEK293T cells transfected with control or two anti-DHP shRNAs (shDHP and shDHP-2) normalized to total DNA. Two independent biological replicates are represented.

